# Chemical inhibition of asexual development leads to increased ultimate tensile strength in mycelial material

**DOI:** 10.1101/2025.05.14.653826

**Authors:** Kelsey Gray, Harley Edwards, Alexander G. Doan, Walker Huso, JungHun Lee, Ethan Folmer, Ololade Lawrence, Mark R. Marten, Steven D. Harris

**Author notes:** 1 Correspondence, 2 Correspondence.

## Abstract

This study investigates the chemical inhibition of asexual development in Aspergillus nidulans to enhance the mechanical properties of mycelial materials. We hypothesized that suppressing conidiation using the ornithine decarboxylase inhibitor α-difluoromethylornithine (DFMO) would increase material strength by inhibiting asexual development, promoting denser hyphal packing. Mycelial materials were grown in DFMO concentrations (0, 0.05, 0.5, and 5 mM), and conidiation and ultimate tensile strength (UTS) were measured. Results showed a dose-dependent reduction in conidiation, with significant decreases at all DFMO levels (P ≤ 0.05). While lower DFMO concentrations (0.05 and 0.5 mM) did not significantly alter UTS, 5 mM DFMO treatment doubled the material’s tensile strength compared to controls (P ≤ 0.05). Scanning electron microscopy confirmed reduced developmental structures in DFMO-treated samples, supporting the hypothesis. The non-linear relationship between conidiation suppression and strength improvement suggests additional mechanisms, such as hyphal morphology or cell wall changes, may contribute. These findings demonstrate that chemical modulation of fungal development can rationally tune mycelial material properties, offering a systematic approach for biomaterial engineering.

## 1. INTRODUCTION

Systematic, rational approaches are now standard practice in modern drug discovery as they reduce knowledge gaps, shorten development cycles, and thus foster innovation (1-8). These systematic approaches enhance understanding of a subject’s capabilities and mechanisms, thus providing rational optimization strategies (9). To advance development of traditional biomaterials, scientists have successfully implemented similar systematic approaches to overcome inherent biological complexities (10-12). In contrast, traditional development of mycelial materials (i.e., sustainable materials composed primarily of filamentous fungi) has employed a “test and see” methodology, adjusting various inputs (e.g., species, nutrients, post-growth processing) to generate diverse observations and deduce material optimization strategies afterwards (13-15). To address this shortcoming, we have attempted to develop a systematic approach for design of mycelial materials. Our working hypothesis has been that genetic changes can be used to rationally tune fungal phenotype, resulting in mycelial materials with specific mechanical properties.

In our initial work, we were surprised to find that deletion of the *A. nidulans* cell-wall repair gene (Δ*mpkA*) leads to mycelial materials with increased tensile strength (16). To determine possible causes, we carried out a comprehensive phenotypic analysis which led us to hypothesize the increased strength was due to lack of asexual developmental structures (e.g., conidiophores), leading to more densely packed hyphae. We tested this hypothesis in a subsequent paper (17) where we generated mycelial material from four previously characterized aconidial mutants. Consistent with our hypothesis, we found that material generated from all four aconidial mutants had greater tensile strength than material generated from a control. Here we sought to expand on this finding, and determine if a chemical approach could be used to repress asexual development resulting in similar increases in tensile strength.

Previous studies have shown that ornithine decarboxylase inhibitors suppress conidiation in fungi (18-20). We hypothesized that growing mycelial materials in the presence of increasing concentrations the ornithine decarboxylase inhibitor, α-difluoromethylornithine (DFMO) would allow us to tune the degree of asexual development. And that this would lead to proportional increases in ultimate tensile strength of the resulting mycelial material.

To test this hypothesis, we measured both degree of conidiation and ultimate tensile strength of mycelial materials generated from the model fungus *Aspergillus nidulans* grown in increasing DFMO concentrations (0, 0.05, 0.5, and 5 mM). Our results show DFMO does reduce asexual development, which leads to an increase in ultimate tensile strength compared to the control of the tested mycelial materials.

## 2. MATERIALS & METHODS

### 2.1 Fungal Strains and Growth Media

The A28 strain (*pabaA6*; *biA1*), acquired from the Fungal Genetic Stock Center (FGSC, Manhattan, KS), was used in all experiments. Initial cultivation was performed on MAG-V agar plates, modified to contain 2 g/L BD Bacto Peptone, 1 mL/L vitamin mix, 1 mL/L Hutner’s trace elements, 20 g/L granulated agar, 1.12 g/L uracil, 1.22 g/L uridine, 35.065 g/L NaCl, 20 g/L glucose, and 20 g/L malt extract (16). The trace element solution consisted of 22 g/L ZnSO□·7H□O, 11 g/L H□BO□, 5 g/L MnCl□·4H□O, 5 g/L FeSO□·7H□O, 1.7 g/L CoCl□·6H□O, 1.6 g/L CuSO□·5H□O, 1.5 g/L Na□MoO□·2H□O, and 50 g/L EDTA (Na□). The vitamin solution included 100 mg/L each of biotin, pyridoxine, thiamine, riboflavin, p-aminobenzoic acid, and nicotinic acid.

For mycelial growth, a modified YGV medium was prepared with 1 g/L yeast extract, 2 g/L BD Bacto peptone, 1 g/L Bacto casamino acids, 1 mL/L vitamin solution, 1 mL/L Hutner’s trace elements, 44.21 g/L KCl, 1.12 g/L uracil, 1.22 g/L uridine, 20 g/L glucose, 20 g/L malt extract, 10 g/L proline, 50 mL/L nitrate salts, and 5 mL/L MgSO□ solution (16). The nitrate salts contained 142.7 g/L KNO□, 10.4 g/L KCl, 16.3 g/L KH□PO□, and 20.9 g/L K□HPO□, while the MgSO□ solution was prepared at 104 g/L. An additional YGV aliquot was supplemented with 1.1835 g/L DFMO (236.7 g/mol).

### 2.2 Growing Mycelial Materials

The A28 strain was cultivated to produce mycelial material using established methods (16). In summary, fungal cultures were grown on modified MAG-V agar plates to produce conidia, which were then collected, quantified, and used to inoculate YGV medium in 60mm Fisherbrand petri dishes. The medium was supplemented with or without DFMO, depending on the experimental group, and incubated at 28°C for 120 hours. After growth, the mycelial disc was harvested and rinsed sequentially with bleach and deionized water.

### 2.3 Asexual Development (Conidiation)

For quantification of conidia, material was grown as above and soaked in hexamethyldisilazane (HMDS) for 5 min then air dried for at least 90 minutes (16). The dried material was added to a cryogenic vial containing 3 glass beads (diameter 3 mm), and homogenized with a Mini Beadbeater (Biospec Products) at 5000 rpm for 5 seconds. Then 1 mL of DI water was added, vortexed for 15 seconds, homogenized again at 5000 rpm for 5 seconds, then vortexed once more for 15 seconds. Homogenized samples were diluted with DI water before being vortexed and placed onto a hemocytometer for counting conidia.

### 2.4 Tensile Testing (Mechanical Testing)

Tensile tests were performed on approximately 4 × 40 mm specimens sectioned (Universal Laser Systems VLS 3.6/6) from mycelial disks. A minimum of five samples were extracted from each disk, with three disks analyzed per genotype. Prior to testing, the specimens were immersed in a 40:60 glycerol-water solution for one minute, lightly coated in petroleum jelly, and briefly dried using warm air for about one minute per side. Exact sample dimensions were determined using a dissection microscope (Leica Microscope s9i Model meb115), after which the specimens were secured to a paper support frame using cyanoacrylate adhesive (Loctite) and NaHCO□ powder (to hasten curing).. The frame was affixed to the tensile tester (Instron 3369) with double-sided tape. Just before testing, the paper frame was carefully trimmed to ensure only the mycelial material underwent tension. Reflective markers were attached to the frame to facilitate strain tracking via a laser extensometer (Electronic Instrument Research Model LE-01). Force data were recorded using a load cell (Transducer Techniques Sensor) integrated with the testing apparatus. Each test was terminated upon sample failure (16).

### 2.5 Scanning Electron Microscopy (SEM)

Scanning electron microscopy (SEM) was performed following established methods (16). Mycelial materials were examined both before and after tensile testing. Samples were mounted on aluminum stubs with double-sided carbon tape, coated with Au/Pd using a Cressington 108 Manual Sputter Coater, and analyzed using an FEI Nova NanoSEM 450.

### 2.6 Significance Testing

All quantitative data was subjected to T-tests to identify any significant differences between groups using Microsoft Excel.

## 3. RESULTS AND DISCUSSION

### 3.1 DFMO Driven Fungal Morphology

To evaluate DFMO’s impact on developmental structures in mycelial material, we quantified conidia production across a concentration gradient (0, 0.05, 0.5, and 5.0 mM DFMO). **Figure 1** demonstrates a significant dose-dependent decrease in conidiation with increasing DFMO concentrations (*P* ≤ 0.05). Statistical analysis revealed significant differences in conidiation between all tested DFMO concentrations (*P* ≤ 0.05 for all pairwise comparisons). Preliminary experiments at concentrations below 0.05 mM and above 5 mM (data not shown) yielded no significant differences compared to their nearest tested concentration, indicating we identified the effective concentration range for DFMO’s inhibitory effects. These results confirm that DFMO treatment progressively reduces asexual developmental structures (conidia) in *A. nidulans* mycelial materials in a concentration-dependent manner.

**Figure 1.**
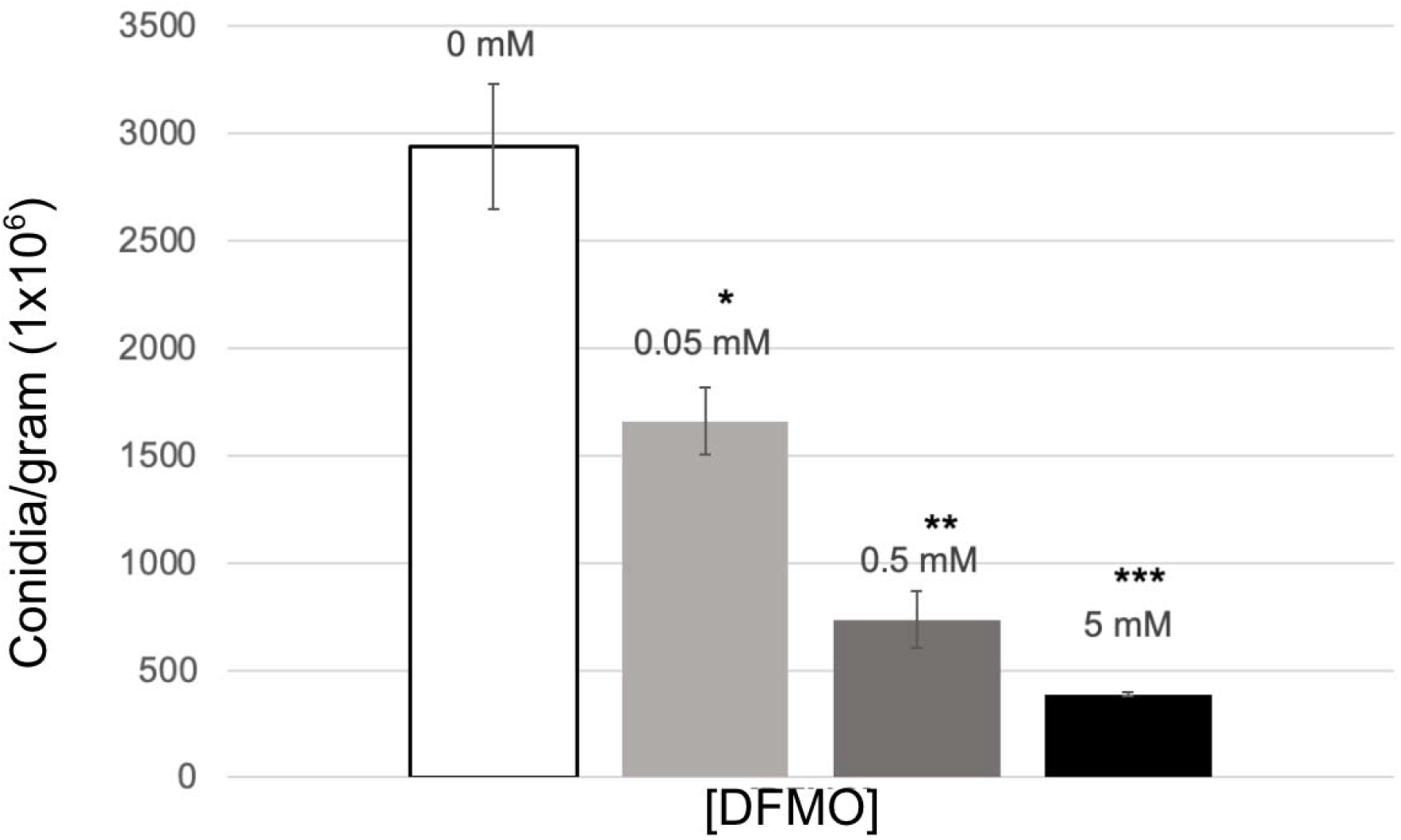
Average number of conidia produced per gram of *Aspergillus nidulans* (A28) mycelial material grown with different concentrations of DFMO. For all bars, *n* = 12; error bars represent standard error. All DFMO concentrations showed significant difference when compared to control. * = *P* ≤ 0.05, 0 mM ** = *P* ≤ 0.05, 0 mM, 0.5 mM *** = *P* ≤ 0.05, 0 mM, 0.5 mM, 5 mM

### 3.2 Mechanical Properties

To assess mechanical properties, mycelial-material discs were prepared according to our established protocol and sectioned into five testing strips. We evaluated three independent discs for each DFMO concentration (0, 0.05, 0.5, and 5.0 mM), yielding 15 test specimens per treatment condition. Tensile testing revealed that all samples exhibited linear elastic behavior before failure, characteristic of brittle materials. Ultimate tensile strength (UTS) analysis, presented in **Figure 2**, demonstrates that materials treated with 0.05 and 0.5 mM DFMO showed increased mean strength values compared to untreated controls, but these differences were not statistically significant. Only mycelial materials treated with 5 mM DFMO exhibited significantly enhanced tensile strength (*P* ≤ 0.05), with approximately twice the UTS of control samples. This concentration-dependent response suggests a threshold effect in DFMO’s ability to improve mechanical performance.

**Figure 2.**
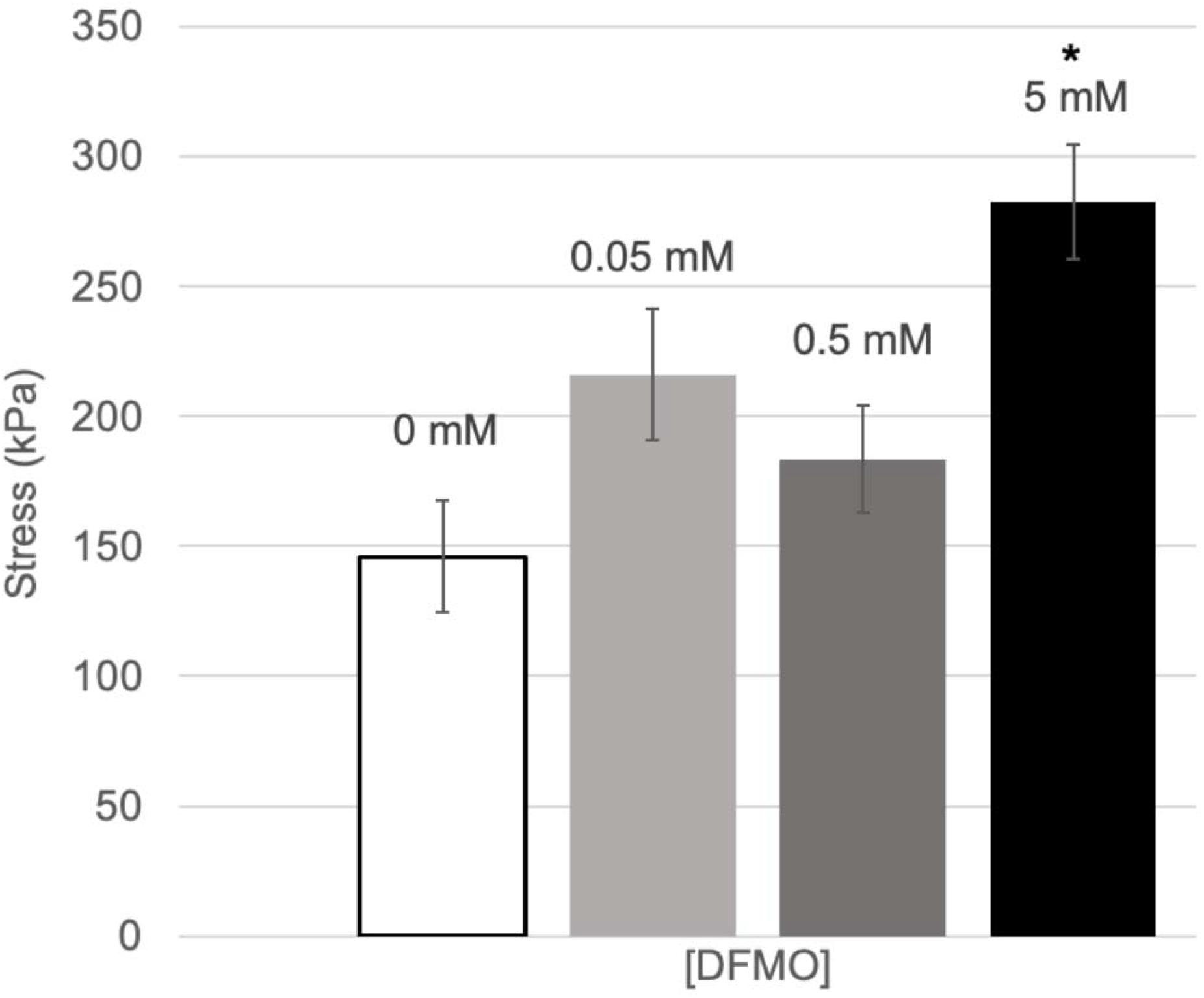
Average ultimate tensile strength (UTS) of *Aspergillus nidulans* (A28) mycelial material grown with different concentrations of DFMO. For all bars, *n* = 15; error bars are standard error; * = *P* ≤ 0.05.

### 3.3 Qualitative Fractional and Surface Based Analysis

To validate our conidiation findings and further characterize the structural basis for altered mechanical properties, we examined fracture surfaces of tensile-tested specimens using scanning electron microscopy (SEM). Representative micrographs presented in **Figure 3** provide compelling qualitative evidence for DFMO-induced suppression of asexual development structures, corroborating our quantitative measurements in **Figure 1**. The progression from control samples (0 mM) to 5 mM DFMO treatment reveals a clear dose-dependent reduction in conidiophores and associated structures at the material surface. This microscopic evidence strengthens our conclusion that DFMO effectively inhibits conidiation in *A. nidulans* mycelial materials and suggests that altered development patterns may contribute to the observed mechanical property improvements.

**Figure 3.**
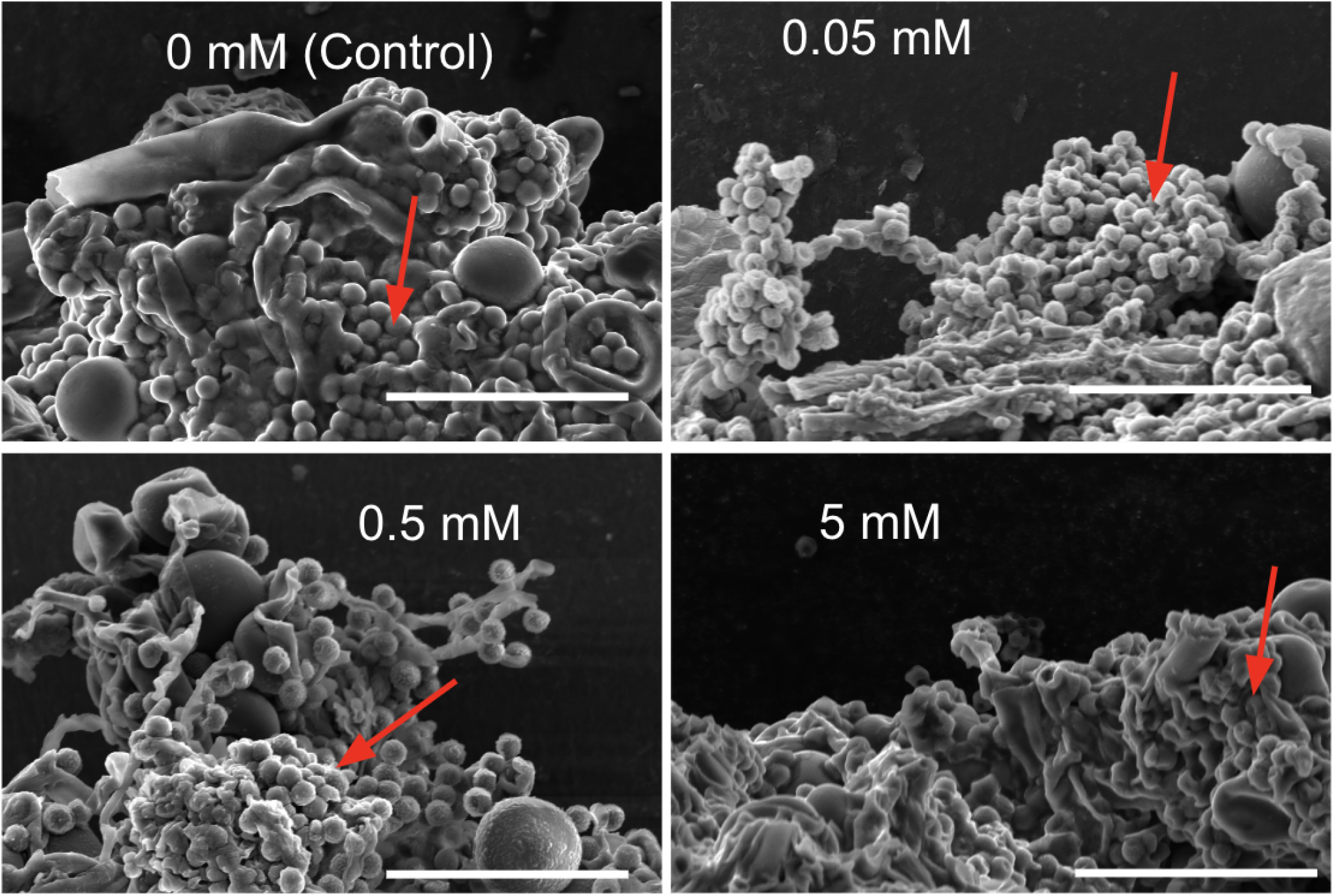
Representative SEM images of *Aspergillus nidulans* (A28) mycelial material showing structure and morphological features at the fracture point after tensile testing. The increasing concentration of DFMO in each image shows a decrease in developmental features. All bars = 30 µm.

## 4. CONCLUSIONS

Our study demonstrates that treatment with 5 mM DFMO significantly increases the ultimate tensile strength of *A. nidulans* mycelial materials while effectively inhibiting asexual development. Interestingly, while conidiation decreases exponentially with increasing DFMO concentration across our treatment range, mechanical strength improvement does not follow the same pattern. This non-linear relationship suggests that the mechanism by which DFMO enhances material properties extends beyond simple conidiation suppression, potentially involving additional alterations to hyphal morphology, cell wall composition, or mycelial network architecture. These findings provide a foundation for developing chemically-modified mycelial materials with enhanced mechanical properties and highlight the potential for DFMO treatment as a tunable strategy for biomaterial engineering.

## Acknowledgments

We gratefully acknowledge significant assistance from Drs. Michael Duffy and Joao Santos (Engineering Testing, LLC) for their assistance and guidance with material tensile testing; Dr. Govind Rao (UMBC, Center for Advanced Sensor Technology) for use of the laser cutter to prepare samples; Dr. Jorge Almodovar for general guidance.

## References

1. Milligan PA, Brown MJ, Marchant B, Martin SW, Van Der Graaf PH, Benson N, et al. Model□based drug development: a rational approach to efficiently accelerate drug development. Clinical Pharmacology & Therapeutics. 2013;93(6):502–14.

2. Huggins DJ, Sherman W, Tidor B. Rational approaches to improving selectivity in drug design. Journal of medicinal chemistry. 2012;55(4):1424–44.

3. Reddy MR, Parrill AL. Overview of rational drug design. ACS Publications; 1999.

4. Mandal S, Moudgil Mn, Mandal SK. Rational drug design. European journal of pharmacology. 2009;625(1-3):90–100.

5. Mavromoustakos T, Durdagi S, Koukoulitsa C, Simcic M, G. Papadopoulos M, Hodoscek M, et al. Strategies in the rational drug design. Current medicinal chemistry. 2011;18(17):2517–30.

6. Ette EI, Garg V, Jayaraj A. A rational approach to drug development: the exploratory phase. Clinical Research and Regulatory Affairs. 2004;21(2):155–77.

7. Stanton RV, Mount J, Miller JL. Combinatorial Library Design:□ Maximizing Model-Fitting Compounds within Matrix Synthesis Constraints. Journal of Chemical Information and Computer Sciences. 2000;40(3):701–5.

8. Lowe G. Combinatorial chemistry. Chemical Society Reviews. 1995;24(5):309–17.

9. Sturm B, Sunyaev A. Design principles for systematic search systems: a holistic synthesis of a rigorous multi-cycle design science research journey. Business & Information Systems Engineering. 2019;61:91–111.

10. Kohn J, Welsh WJ, Knight D. A new approach to the rationale discovery of polymeric biomaterials. Biomaterials. 2007;28(29):4171–7.

11. Gao H. Modelling strategies for nano-and biomaterials. European White Book on Fundamental Research in Materials Science Stuttgart (Germany): Max Planck Gesellschaft, GMBH. 2001:144–8.

12. Brocchini S, James K, Tangpasuthadol V, Kohn J. Structure–property correlations in a combinatorial library of degradable biomaterials. Journal of Biomedical Materials Research: An Official Journal of The Society for Biomaterials, The Japanese Society for Biomaterials, and the Australian Society for Biomaterials. 1998;42(1):66–75.

13. Haneef M, Ceseracciu L, Canale C, Bayer IS, Heredia-Guerrero JA, Athanassiou A. Advanced materials from fungal mycelium: fabrication and tuning of physical properties. Scientific reports. 2017;7(1):41292.

14. Deeg K, Gima Z, Smith A, Stoica O, Tran K. Greener Solutions: Improving performance of mycelium-based leather. Final Report to MycoWorks. 2017:1–24.

15. Appels FV, van den Brandhof JG, Dijksterhuis J, de Kort GW, Wösten HA. Fungal mycelium classified in different material families based on glycerol treatment. Communications biology. 2020;3(1):1–5.

16. Gray K, Edwards H, Doan AG, Huso W, Lee J, Pan W, et al. Aspergillus nidulans cell wall integrity kinase, MpkA, impacts cellular phenotypes that alter mycelial-material mechanical properties. Fungal Biology and Biotechnology. 2024;11(1):22.

17. Gray KJ, Edwards H, Doan AG, Huso W, Lee J, Pan W, et al. The impact of fungal developmental structures on mechanical properties of mycelial materials. bioRxiv. 2025:2025–04.

18. Guzmán-de-Peña D, Aguirre J, Ruiz-Herrera J. Correlation between the regulation of sterigmatocystin biosynthesis and asexual and sexual sporulation in Emericella nidulans. Antonie Van Leeuwenhoek. 1998;73:199–205.

19. Birecka H, Garraway MO, Baumann RJ, McCann PP. Inhibition of ornithine decarboxylase and growth of the fungus Helminthosporium maydis. Plant Physiology. 1986;80(3):798–800.

20. Valdés-Santiago L, Cervantes-Chávez JA, León-Ramírez CG, Ruiz-Herrera J. Polyamine metabolism in fungi with emphasis on phytopathogenic species. Journal of Amino Acids. 2012;2012(1):837932.

